# Processing-bias correction with DEBIAS-M improves cross-study generalization of microbiome-based prediction models

**DOI:** 10.1101/2024.02.09.579716

**Authors:** George I. Austin, Aya Brown Kav, Heekuk Park, Jana Biermann, Anne-Catrin Uhlemann, Tal Korem

## Abstract

Every step in common microbiome profiling protocols has variable efficiency for each microbe. For example, different DNA extraction kits may have different efficiency for Gram-positive and -negative bacteria. These variable efficiencies, combined with technical variation, create strong processing biases, which impede the identification of signals that are reproducible across studies and the development of generalizable and biologically interpretable prediction models. “Batch-correction” methods have been used to alleviate these issues computationally with some success. However, many make strong parametric assumptions which do not necessarily apply to microbiome data or processing biases, or require the use of an outcome variable, which risks overfitting. Lastly and importantly, existing transformations used to correct microbiome data are largely non-interpretable, and could, for example, introduce values to features that were initially mostly zeros. Altogether, processing bias currently compromises our ability to glean robust and generalizable biological insights from microbiome data. Here, we present DEBIAS-M (**D**omain adaptation with phenotype **E**stimation and **B**atch **I**ntegration **A**cross **S**tudies of the **M**icrobiome), an interpretable framework for inference and correction of processing bias, which facilitates domain adaptation in microbiome studies. DEBIAS-M learns bias-correction factors for each microbe in each batch that simultaneously minimize batch effects and maximize cross-study associations with phenotypes. Using benchmarks of HIV and colorectal cancer classification from gut microbiome data, and cervical neoplasia prediction from cervical microbiome data, we demonstrate that DEBIAS-M outperforms batch-correction methods commonly used in the field. Notably, we show that the inferred bias-correction factors are stable, interpretable, and strongly associated with specific experimental protocols. Overall, we show that DEBIAS-M allows for better modeling of microbiome data and identification of interpretable signals that are reproducible across studies.

## Introduction

A hallmark of a robust scientific analysis is that its conclusions generalize beyond a specific processing protocol, study, or population. Such generalization offers strong evidence that the findings are not the result of the particularities of one experiment, reduces the impact of confounding variables, and, in general, lowers the risk for spurious findings. For prediction models, external validation in an independent study is imperative for a robust assessment of the generalizability of the model to new populations^1,2^. The ability to train generalizable models across datasets also offers an opportunity for increased sample size and power in settings where data from many smaller studies is already available, such as studies of the vaginal microbiome in preterm birth^3–8^, or the gut microbiome in colorectal cancer^9,10^.

In the modern era of sequencing-based culture-independent microbiome profiling, challenges in generalizability stem not only from biological variability, such as differences between populations, study design, and medical or cultural practices, but also from substantial variability between microbiome profiling protocols, facilities, and bioinformatic analysis pipelines. Such variability has been noted early and repeatedly^11–17^, and substantially affects the replicability and interpretability of microbiome data^18^. Large-scale comparative efforts conducted across different laboratories concluded that variation between protocols could even surpass biological variation^19,20^. While this has prompted calls for standardization of protocols across the field, others doubt the utility of this strategy^21^, and, in practice, wide adoption of uniform and standardized protocols has not taken place.

In a landmark study, McLaren, Willis & Callahan provided and validated a mathematical theory and foundation for consideration of variability in microbiome studies^22^. Under their framework, technical variability manifests as taxon-specific multiplicative bias that stems from differential efficiencies at different steps of the experimental and analytic pipeline, including DNA extraction, PCR, sequencing, and bioinformatics analyses. While DNA extraction was highlighted as a significant source of bias^19,22^, McLaren et al. demonstrate that each step has different efficiencies for each taxon, together acting to distort the measured microbial abundances. As each taxon and every study have their own unique biases, statistically significant interactions (e.g., with a phenotype) measured in one study might not replicate in a second one (**Fig. 1a**). Even if individual studies separately show similar signals, biases can also produce different signals on their aggregation due to distribution shifts (known as Simpson’s paradox). Importantly, McLaren et al. show that even if identical taxon-specific biases apply across samples, their effect on each sample depends on its microbial composition, and is therefore not eliminated even with rigorous standardization of experimental factors^22^. Hence, there is a pressing need for methods that can retrospectively infer and correct processing biases introduced during microbiome profiling.

**Figure 1.**
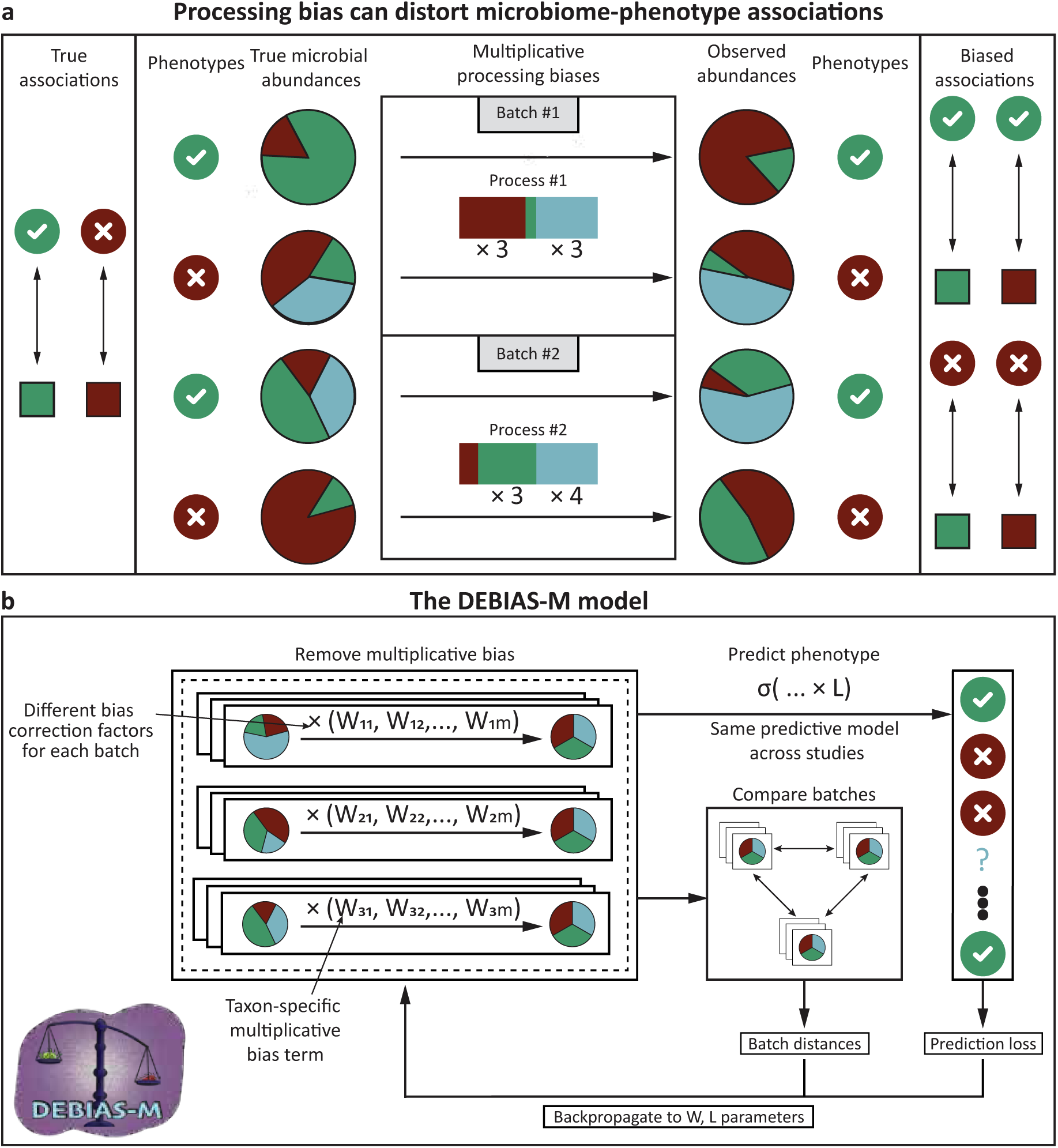
Processing-bias correction with DEBIAS-M. **a,** An illustration of our assumptions underlying the data-generating process: The “real” relative abundances of a given sample are distorted by batch-specific per-taxa multiplicative biases, which can both obfuscate the true signals or generate artifactual ones, particularly in cross-batch analyses. **b,** DEBIAS-M iteratively learns per-species multiplicative biases for each batch which minimize the distances between the batches and maximize linear associations to a phenotype of interest. Samples without available phenotype (missing or hidden) are used only when calculating the batch-wise differences.

Many approaches relevant to this challenge have been developed under the general term of batch-correction methodologies, some specifically developed for microbiome data^23–28^ and others adopted from other fields^29–32^. While these methods show some benefit in correcting batch effects, many of them make strong parametric assumptions that do not necessarily apply to microbiome data, which are sparse and zero-inflated. Other methods are suitable only for association testing, rather than data correction and predictive modeling. Furthermore, some methods require the use of an outcome variable, which risks overfitting the data and limits the ability to test for external generalization. Lastly and importantly, the changes these methods make to the data are largely non-interpretable, as recently highlighted with respect to the application of voom-SNM^30,31,33^ to microbial reads identified in tumor sequencing data, in which values were introduced to features that were initially very sparse^33,34^.

In this work, we present “Domain adaptation with phenotype Estimation and Batch Integration Across Studies of the Microbiome” (DEBIAS-M). DEBIAS-M is a method for inference and correction of processing bias in microbiome data within a phenotype prediction framework, designed to operate in the context of multiple processing batches or studies (collectively termed “batches”). DEBIAS-M infers taxon- and batch-specific processing biases by finding batch-wise patterns, which, when corrected for, both reduce differences between studies and improve the overall association with phenotypes of interest. It does so while accounting for the compositional nature of microbiome data, which enables the method to integrate effectively within standard microbiome analysis frameworks. We demonstrate that DEBIAS-M outperforms batch-correction methods using diverse benchmarks of HIV, colorectal cancer, and cervical neoplasia predictions across gut and vaginal microbiome studies, including the use of both metagenomics and 16S sequencing. We further show that the bias-correction factors learned by DEBIAS-M are interpretable and strongly associated with experimental protocols. Finally, we demonstrate that incorporating DEBIAS-M into common machine learning pipelines improves prediction accuracy, and demonstrate generalizable cross-study prediction of cervical neoplasia using vaginal microbiome data. Overall, DEBIAS-M offers the capability to leverage more datasets for explainable, stronger, and more replicable microbiome analyses by minimizing the impact of processing bias.

## Results

### Description of DEBIAS-M

DEBIAS-M is a method for *in silico* detection and correction of processing bias. It takes as input a representation (e.g., read count or relative abundances of taxa) of microbiome samples from multiple processing protocols, studies or batches (hereafter collectively termed “batches”), and learns one multiplicative coefficient^22^ for each taxon in every batch, which corrects for the experimental bias of that batch (**Fig. 1a**). Every sample is then renormalized before a downstream prediction model – identical across all batches – learns an association to a phenotype of interest (**Fig. 1b**; **Methods**). The bias and prediction parameters are optimized to maximize the prediction likelihood while minimizing the domain shifts between batches. Samples for which the phenotype is unavailable or hidden, such as those with missing data or samples in a test set, are not included in the calculation of the prediction loss, and they are only considered when minimizing cross-batch differences (**Fig. 1b**). DEBIAS-M is available as a python package from https://github.com/korem-lab/DEBIAS-M, with a complete description available in the **Methods** section.

### DEBIAS-M outperforms batch-correction methods in diverse benchmarks

We first aimed to evaluate the bias-correction capabilities of DEBIAS-M and compare them to popular batch-correction approaches. While many batch-correction methods are available and have been used in microbiome analyses, we chose to focus on three methods that are either popular or recently developed: ComBat^29^, voom-SNM^30,31,33^, and ConQuR^23^, in addition to using the raw data with no batch correction. To this end, we set up three cross-study prediction benchmarks in which we applied each batch-correction method followed by a classifier of a relevant outcome (**Methods**). For each prediction task, we used a leave-one-study-out approach, where a model was trained on all studies except one, and evaluated on the hidden study. The underlying assumption of this benchmark is that improved batch correction should result in improved classification accuracy on held-out studies. To focus this benchmark on batch correction rather than elaborate machine-learning pipelines, we used linear models. No outcome labels from the hidden study were provided during batch correction or model fitting.

First, we analyzed a benchmark used in previous batch-correction studies^23^, in which gut 16S rRNA gene amplicon sequencing data collected from 17 case-control studies with publicly available data was used to classify HIV diagnosis^35^ (N of 13-233 subjects per study, for a total of 1,032 subjects; **Methods, Table S1**). We found that DEBIAS-M outperformed raw data and data corrected by ComBat, ConQuR and Voom-SNM (median [IQR] auROCs of 0.7 [0.61-0.75] vs. 0.5 [0.45-0.53], 0.53 [0.49-0.58], 0.54 [0.49-0.57], and 0.53 [0.51-0.63], respectively; Fisher’s multiple comparison of DeLong tests *p*<0.01 for all pairwise comparisons with alternative methods; **Fig. 2a**).

**Figure 2.**
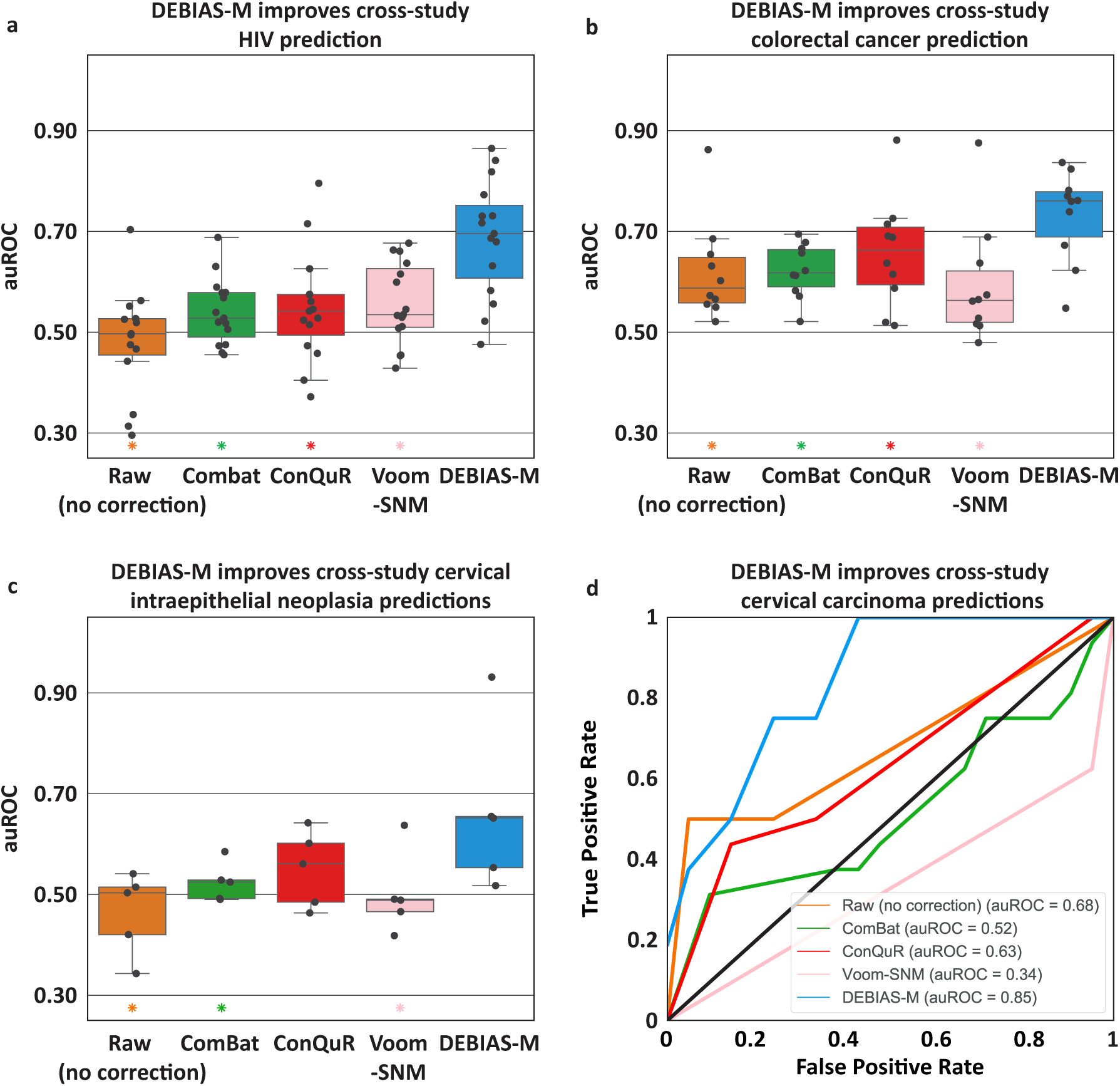
DEBIAS-M outperforms batch-correction methods in cross-study prediction benchmarks. **a-c,** Box and swarm plots (Box, IQR; line, median; whiskers, nearest point to 1.5*IQR) of auROCs, each evaluating the generalization performance of logistic regression models to a held-out study, predicting: HIV from gut microbiome data (**a**; N=13-233 samples per study; **Methods**); colorectal cancer from gut microbiome data (**b**; N=53-128 samples per study); and cervical intraepithelial neoplasia from cervical microbiome data (**c**; N=29-82 samples per study). **d,** ROC plot of logistic regression models predicting cervical carcinoma in a held-out study using cervical microbiome data (N=293 for 4 studies in the training set and N=29 samples in the held-out study). * denotes *p<*0.01 for DEBIAS-M vs marked method, via Fisher’s multiple comparison of DeLong tests.

We then performed a similar cross-study evaluation of predictions using publicly available data from metagenomic sequencing data of gut microbiome samples in studies evaluating colorectal cancer^9,10,36–41^. In this analysis, we once again saw that DEBIAS-M results outperformed alternative methods, with raw data, ComBat, ConQuR and voom-SNM yielding median [IQR] auROCs of 0.58 [0.57-0.65], 0.62 [0.59-0.66], 0.66 [0.59-0.71], and 0.56 [0.52-0.62], compared to 0.76 [0.69-0.78] for DEBIAS-M (*p*<0.01 for all; **Fig. 2b**).

Next, we performed a similar benchmark using the cervical microbiome to predict cervical neoplasia, a challenging scenario with potential translational implications. We therefore compiled and uniformly analyzed data from 5 independent studies that recruited patients with and without cervical intraepithelial neoplasia and cervical cancer^42–46^ (N of 29-82 subjects per study, for a total of 322 subjects; **Methods**, **Table S2**). First, we used these datasets to do a cross-study evaluation of classification of cervical intraepithelial neoplasia, a precancerous state conveying high risk for cervical cancer. While the linear models trained on raw data and data corrected by ComBat, ConQuR, and voom-SNM yielded median [IQR] auROCs of 0.50 [0.42-0.51], 0.52 [0.49-0.53], 0.56 [0.48-0.6], and 0.49 [0.47-0.49], respectively, DEBIAS-M had cross-study auROCs of 0.65 [0.55-0.65] (DeLong p<0.01 for all pairwise comparisons with DEBIAS-M, except with ConQuR, *p*=0.048; **Fig. 2c**). We then devised models for classifying the presence of cervical carcinoma, for which phenotypes were available only for two studies^42,45^. Using the smaller of the two studies as a test set, we found that DEBIAS-M was substantially more accurate than all other methods (auROC=0.85 vs. 0.34-0.68, p=0.02, 0.07, 2.3×10^−4^ and 0.25 for ComBat, ConQuR, Voom-SNM, and the raw data, respectively; **Fig. 2d**). Overall, DEBIAS-M demonstrates robust performance improvement compared with batch-correction methods, demonstrated across three different benchmarks totaling 2,240 samples from 32 studies, including both vaginal and gut microbiome, 16S and metagenomic sequencing.

### DEBIAS-M is robust for training and testing strategy and for operating in log space

Multiple strategies have been used in previous studies^23,26,33,47,48^ with respect to using features (i.e., microbiome data) and outcomes (i.e., phenotypes) during batch correction, which impacts the evaluation of machine learning models (**Fig. S1**). We have used a common approach, in which microbiome and covariates data are available for all samples (from both the training and test sets) during batch correction, but the phenotype labels are only available for samples from the training set (**Fig. S1c**), ensuring that there is no “information leakage” from the test set^47^. We note, however, that in some previous studies batch-correction methods have incorporated phenotype labels from all samples during batch-correction (including the test set), thus allowing for information leakage (**Fig. S1b**), and that doing so in our benchmarks drastically inflates predictive performance (**Fig. S2**). Additionally, as some concerns have been raised regarding the use of features and covariates from the test set during data processing^47^, we also implemented a variation of DEBIAS-M that performs batch correction and model training using only the training set, and adapts the inferred correction factors on the test set only once the rest of the model parameters are frozen (**Methods**; **Fig. S1d**). DEBIAS-M performs equivalently both when observing test-set features and covariates (**Fig. S1c**) and when these remain “hidden” (**Fig. S1d**; **Fig. S3**), demonstrating that it is robust to training and evaluation strategy.

While many batch-correction methods operate in count or relative abundance space, and so does DEBIAS-M, the use of compositional transformations, such as the centered log-ratio transformation (clr), is often recommended. We therefore performed benchmarks in which DEBIAS-M performed log-additive correction (**Methods**), and was compared with data provided by batch-correction methods after clr transformation. DEBIAS-M outperformed batch-correction methods in all three datasets also in this benchmark, demonstrating its robustness to operating in log space (DeLong *p*<0.01 for all but one pairwise comparisons of ConQuR with DEBIAS-M; **Fig. S4**). We note that the additive property of biases in log-space is likely one reason explaining why prediction models based on microbiome data might be often more effective in this space, as multiplicative biases will have a weaker effect on observed associations in log space.

### DEBIAS-M is robust to dataset characteristics

To evaluate the performance of DEBIAS-M against a known ground truth, we created synthetic simulations of multiple microbiome datasets under different processing biases. We used data-driven simulations using an established microbiome data generator^49^ that we trained on real microbiome data^50^. We then used the bias framework proposed by McLaren et al.^22^ to distort the ground-truth relative abundances with multiplicative species-specific bias terms. We generated synthetic phenotype labels with varying association strength with the ground-truth (“pre-biased”) microbial abundances (**Methods**). After providing DEBIAS-M with the biased samples and the phenotypes, we compared the Jensen-Shannon divergence between the ground-truth pre-biased abundances and the output from DEBIAS-M; the same comparison was also done for the uncorrected abundances.

Analyzing the effect of different factors on the performance of DEBIAS-M, we found that it showed similar performance with variable sequencing depths (1,000-100,000 reads per sample, **Fig. 3a**) and strength of association with phenotype (weak to strong associations; **Methods**; **Fig. 3b**). However, we have noted an improvement in performance with larger batch size (24-96 samples per batch), particularly when batch size was larger than 24 samples (**Fig. 3c**). We have also found a slight improvement when more batches were available, particularly with more than two batches (**Fig. 3d**). Finally, while a smaller feature space has led to a better performance of DEBIAS-M (median Jensen-Shannon divergence from ground truth of 0.04 and 0.08 for 100 and 1,000 features, respectively), the improvement compared to uncorrected abundances was consistent (**Fig. 3e**). Importantly, and despite the small variations in results across certain parameters, bias correction with DEBIAS-M produced microbial abundances that were more similar to the underlying ground truth than those not corrected by DEBIAS-M across all ranges of simulated parameters (**Fig. 3**, one-sided Wilcoxon signed-rank *p*<0.001 for all comparisons). Overall, these simulation results demonstrate that DEBIAS-M’s performance is robust, suggesting that it could be used on datasets with a wide range of characteristics.

**Figure 3.**
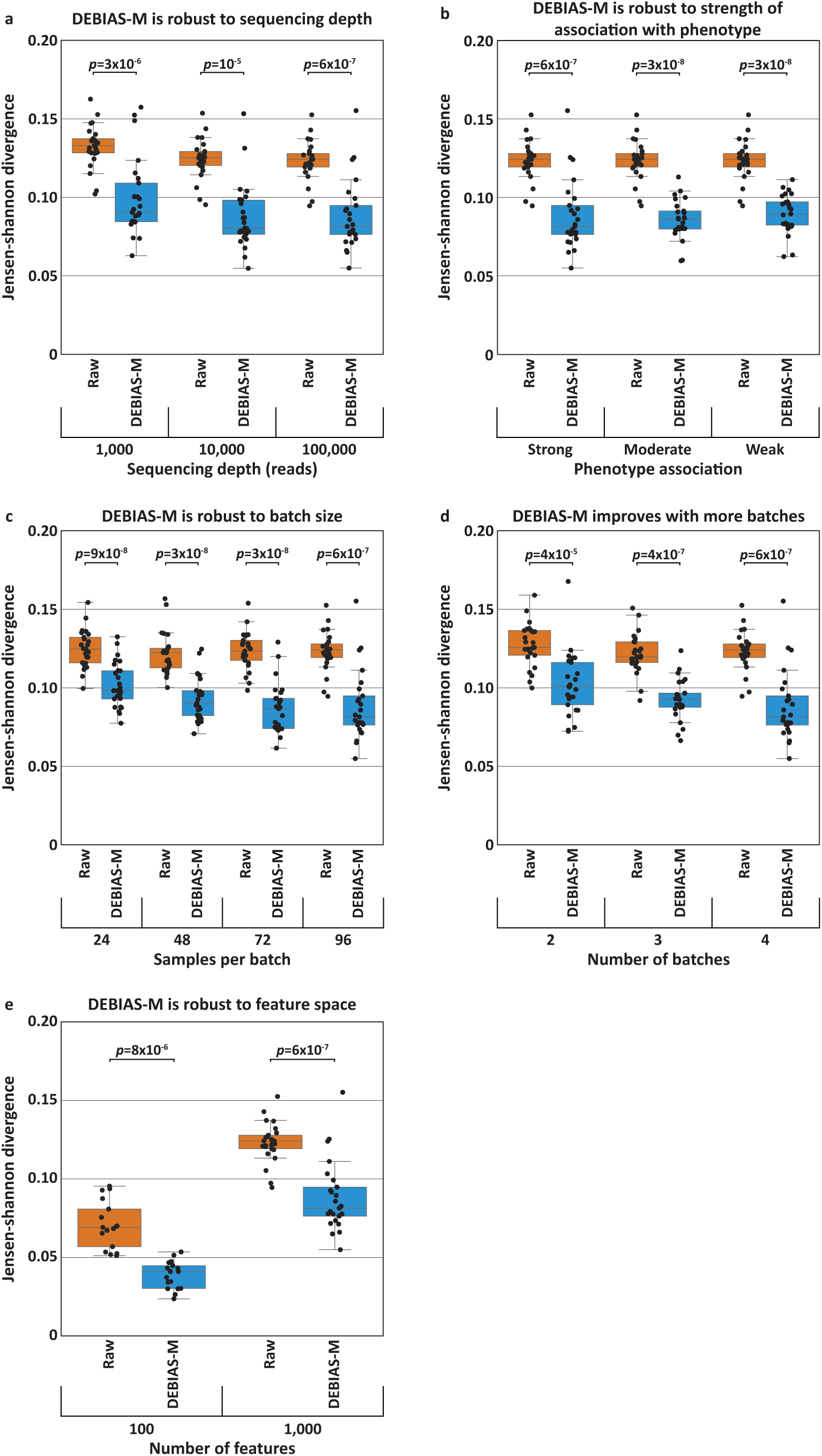
DEBIAS-M is robust across dataset characteristics. Box and swarm plots (line, median; box, IQR; whiskers, nearest point to 1.5*IQR), showing results for *in silico* simulations built under our generative model. DEBIAS-M consistently brings the relative abundance of samples closer to the known ground truths under varying sequencing depth (**a**), strength of association with phenotype (**b**), batch size (**c**), number of batches (**d**), and number of microbiome features (**e**). N=25 experiments per box. In all cases, DEBIAS-M improves the representation of the simulated microbiomes. *p,* Wilcoxon signed-rank test.

### DEBIAS-M learns interpretable bias-correction factors

The interpretability of batch-correction methods came into the spotlight recently^33,34^. Understanding what drives the changes such methods make to the data informs a better interpretation of the analysis as a whole. We therefore next wished to evaluate whether the batch correction performed by DEBIAS-M is interpretable, and whether the correction factors learned for each study could be assigned biological meaning. First, we evaluated whether the correction factors inferred for each batch are stable, which would be expected if they are mostly driven by experimental conditions that are consistent for every batch. Analyzing the same HIV dataset, we applied DEBIAS-M to samples and labels from all studies except a randomly selected half of one held-out study. We then repeated the same procedure for the complementary half of the held-out study, and compared the bias-correction factors learned by DEBIAS-M for each half. We found that these correction factors were highly consistent across halves of different studies (Pearson R=0.59, *p*<0.001, **Fig. 4a**), demonstrating that DEBIAS-M learns consistent bias-correction factors and supporting our hypothesis that these are grounded in processing biases.

**Figure 4.**
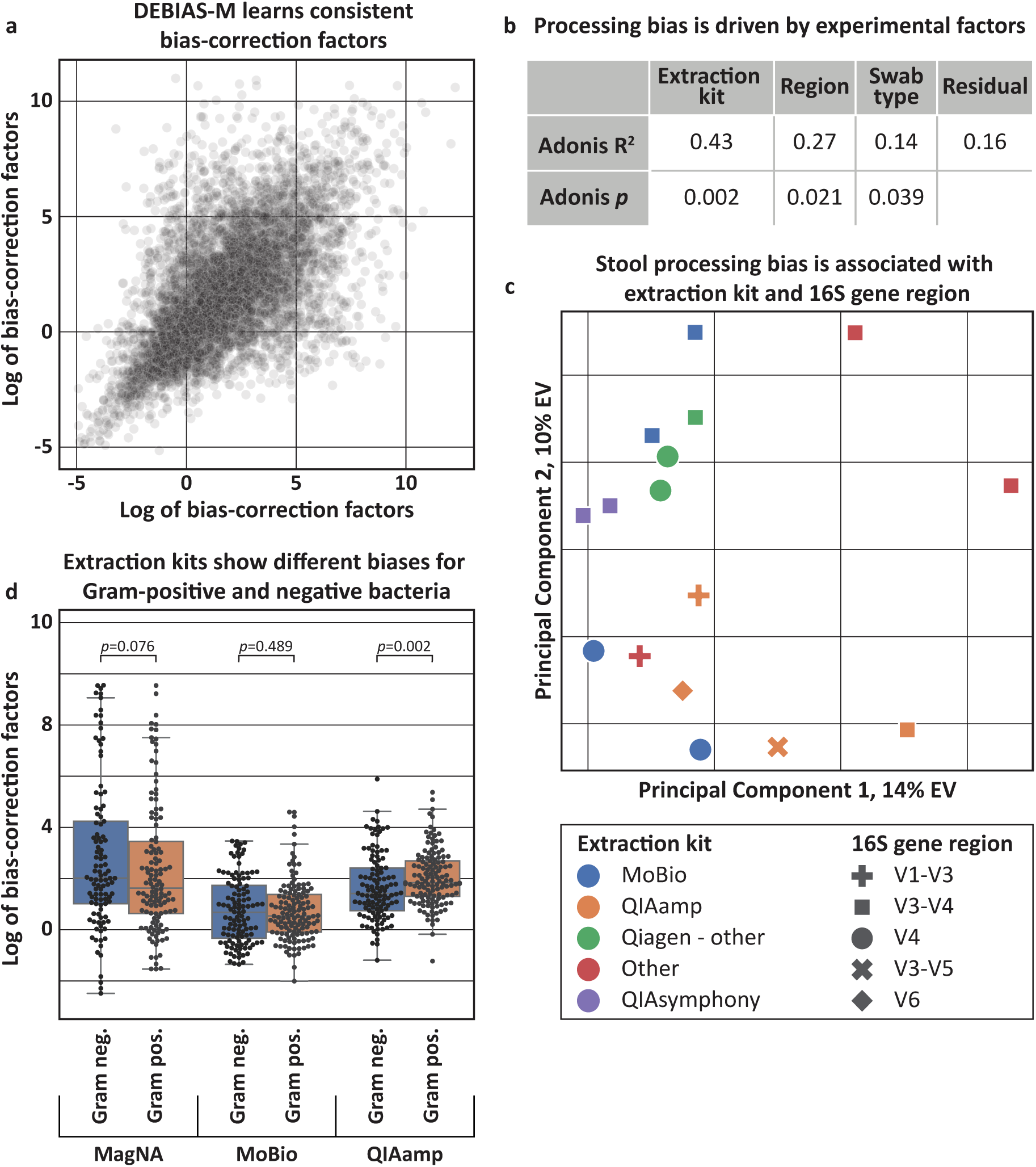
DEBIAS-M infers biases that are stable, interpretable, and consistent with experimental processes. **a,** Scatterplot of bias-correction factors inferred by DEBIAS-M separately for two halves of the same held-out study, evaluated across all studies from the HIV benchmark. Inferred biases are consistent for the same batch (Pearson R=0.59; *p*<0.001). **b,** Adonis PERMANOVA explained variance and p-values for the association of different experimental factors (**Table S1**) with the bias-correction factors inferred by DEBIAS-M for each HIV study. Extraction kit type has the strongest effect on bias. **c,** PCA plot of the bias-correction factors inferred by DEBIAS-M. Color represents the extraction kit type and shape the 16S rRNA region used. **d,** Box and swarm plots (line, median; box, IQR; whiskers, nearest point to 1.5*IQR) showing the bias-correction factors inferred by DEBIAS-M (y-axis) for taxa from the HIV benchmark, stratified by sequencing kit and Gram status. *p*, Mann-Whitney U test.

Next, we sought to find which experimental processing properties are associated with the bias-correction factors inferred by DEBIAS-M. We therefore curated technical parameters for every study in the HIV dataset (**Table S1**) and analyzed them with respect to the bias-correction factors inferred by DEBIAS-M (used in **Fig. 2a**). We found that DNA extraction kit was the most important factor driving the biases learned by DEBIAS-M, accounting for 43% of variance (Adonis^51^ PERMANOVA *p*=0.002; **Fig. 4b,c**), in line with results from a previous analysis of a different dataset^22^. We further found strong associations between the learned bias-correction factors and both 16S region and sample type (fecal, rectal swab, etc.), explaining an additional 27 and 14% of the variance in bias-correction factors, respectively (*p*=0.021 and *p*=0.039, respectively; **Fig. 4b,c**). While the type of extraction kit used was most strongly associated with the learned bias-correction factors, we found that the detection (i.e., presence or absence) of certain bacteria from different studies is instead associated with the choice of 16S region (**Fig. S5a-c**), as previously described^52–54^.

Finally, we investigated whether there are taxon-specific factors that are associated with the inferred processing bias. Comparing the microbe-specific bias-correction factors of Gram-positive and negative bacteria, we found a significant difference between the biases in samples processed with QIAamp kits (Mann-Whitney *U p=*0.002, 0.076, 0.489 for QIAamp, MagNA, and MoBio, respectively; **Fig. 4d**). Such variability between microbiome experimental protocols was previously demonstrated to be associated with Gram status in highly standardized settings^19^. We further noted higher variance in bias-correction factors for studies that used robotic rather than manual experimental processes (Mann-Whitney U *p*=0.03; **Fig. S5d**). DEBIAS-M’s ability to detect microbe- and experimental-specific factors, in a kit-specific fashion and across highly variable studies with a large number of additional processing confounders, demonstrates the sensitivity of our approach. We note that other taxon-specific attributes, such as 16S copy number, had a substantially weaker association with bias-correction factors (Pearson R = -0.10, *p=*0.024; **Fig. S5e**). Altogether, our results demonstrate the interpretability of the bias-correction factors inferred by DEBIAS-M and indicate that they potentially reflect genuine biological factors driving differences between processing protocols.

### Multi-task learning with DEBIAS-M improves metabolite predictions

Because the bias-correction factors inferred by DEBIAS-M are associated with experimental design, a single set of correction factors should generalize for prediction of multiple phenotype labels. To test this hypothesis, we next evaluated DEBIAS-M in a multi-task setting, in which a single set of bias-correction factors is learned per study alongside multiple models that predict different phenotypes (**Fig. 5a**; **Methods**). We therefore designed a benchmark in which the same vaginal microbiome data is used to predict the levels of multiple metabolites measured from paired samples collected simultaneously^3,55^. This is a challenging task, as many metabolites are likely to be affected by factors such as the host or environmental exposures^3,56^. Therefore, we benchmarked the accuracy of models using bias-corrected microbial features to classify whether each of 509 different metabolites had relatively high abundances, evaluating the generalization of these models from one microbiome processing batch to another (**Methods**). First, we used similar benchmarks as above to compare the single-task version of DEBIAS-M to batch-correction methods. Predictions made using DEBIAS-M were significantly more accurate, with a median [IQR] cross-batch auROC of 0.67 [0.6-0.75] across 509 metabolites, compared to 0.57 [0.5-0.64], 0.57 [0.51-0.64], and 0.55 [0.49-0.62] for ComBat, ConQuR, and no correction, respectively (one-sided Wilcoxon *p<*0.001 for all comparisons; **Fig. 5b**).

**Figure 5.**
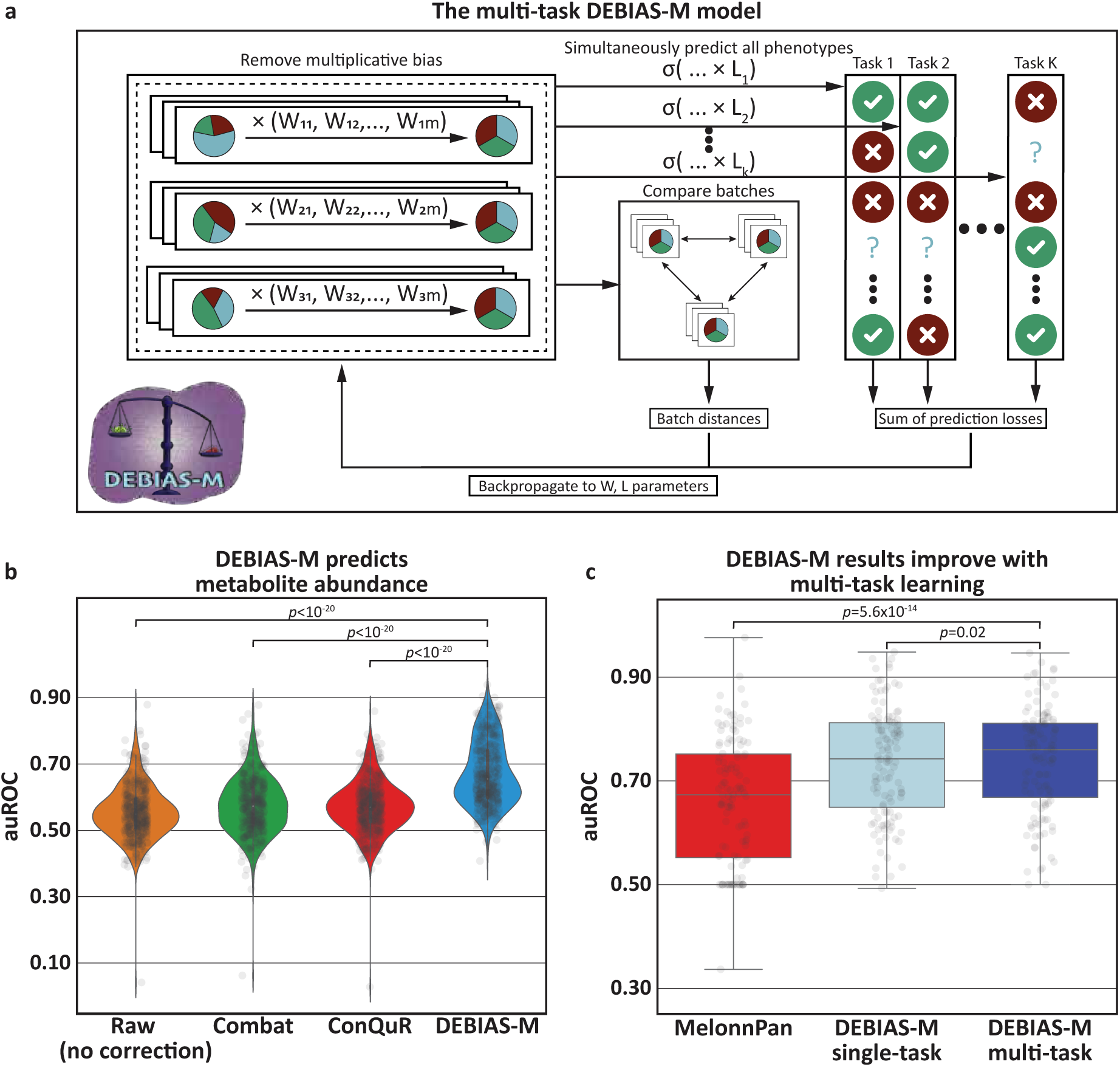
Multi-task learning with DEBIAS-M improves metabolite predictions. **a,** Description of the multi-task version of DEBIAS-M, which learns a single set of bias-correction factors per study while jointly considering multiple prediction tasks. **b,** Violin plot of auROCs from logistic regression models predicting high abundance of 509 metabolites separately, evaluated for cross-batch generalization. **c,** Box and swarm plots (box, IQR; line, median; whiskers, nearest point to 1.5*IQR) of similar cross-batch auROCs for 120 metabolites selected by MelonnPan. The multitask version of DEBIAS-M outperformed both its single-task version (*p*=0.02) and MelonnPan (*p*=5.6×10^−14^). *p*, one-sided Wilcoxon signed-rank test.

Next, we implemented a multi-task version of DEBIAS-M (**Methods**), and compared it both to the single-task version and to MelonnPan^57^, a method for predicting metabolite level from microbiome data. We retrained MelonnPan with default parameters and without any batch correction, and evaluated all three methods on 120 metabolites selected by MelonnPan (**Methods**). The multi-task version of DEBIAS-M outperformed both the single-task DEBIAS-M and MelonnPan, with a median [IQR] auROC of 0.76 [0.67-0.81], compared to a median [IQR] auROC of 0.74 [0.65-0.81] and 0.67 [0.55-0.75], respectively (Wilcoxon *p*=0.02 and *p*=5.6×10^=14^, respectively; **Fig. 5c**). This improvement in predictive performance suggests once again that the bias-correction factors learned by DEBIAS-M are not task-specific, and illustrates the potential of transfer learning in microbiome models in general.

### Preprocessing with DEBIAS-M improves prediction generalizability across cancer studies

Having favorably compared DEBIAS-M to other batch-correction methods, we sought to evaluate its utility in practical scientific investigations, which would often employ more expressive machine-learning algorithms. To this end, we reran our cross-cohort analysis associating the cervix microbiome with cervical neoplasia (**Fig. 2c,d**), except that this time we employed DEBIAS-M as a preprocessing step, whose output was then used to train a random forest model with hyperparameter tuning (**Methods**). For prediction of cervical carcinoma, the random forest model did not improve on results from the linear model (auROCs of 0.63 and 0.68 with and without random forest, respectively;, Delong *p*=0.4; auPRs of 0.45 and 0.63; **Fig. 6a,b**). Preprocessing with DEBIAS-M improved the predictive performance of both, but again with no significant advantage for using random forest models (auROC of 0.84 and 0.85 for DEBIAS-M preprocessing with and without random forest models, *p*=0.47; auPR of 0.68 and 0.71; **Fig. 6a,b**). The same was observed for prediction of cervical intraepithelial neoplasia. While a random forest model offered some improvement over a linear model (median [IQR] auROC of 0.52 [0.48-0.52] and 0.50 [0.42-0.51];, Wilcoxon *p*=0.16), using DEBIAS-M improved on both (0.65 [0.56-0.66] and 0.65 [0.55-0.65] with and without random forest, respectively; *p*=5.6×10^−5^ for comparing random forest models with and without DEBIAS-M; **Fig. 6c**). We note that DEBIAS-M increases the measured α diversity of cervical microbiome samples (Wilcoxon *p*<10^−20^, **Fig. 6d**), although it maintained the same presence and absence of all taxa (**Fig. 6e**). These results demonstrate that bias removal may have a stronger effect on model generalization in a cross-study prediction setting than the selection of a particular machine learning algorithm.

**Figure 6.**
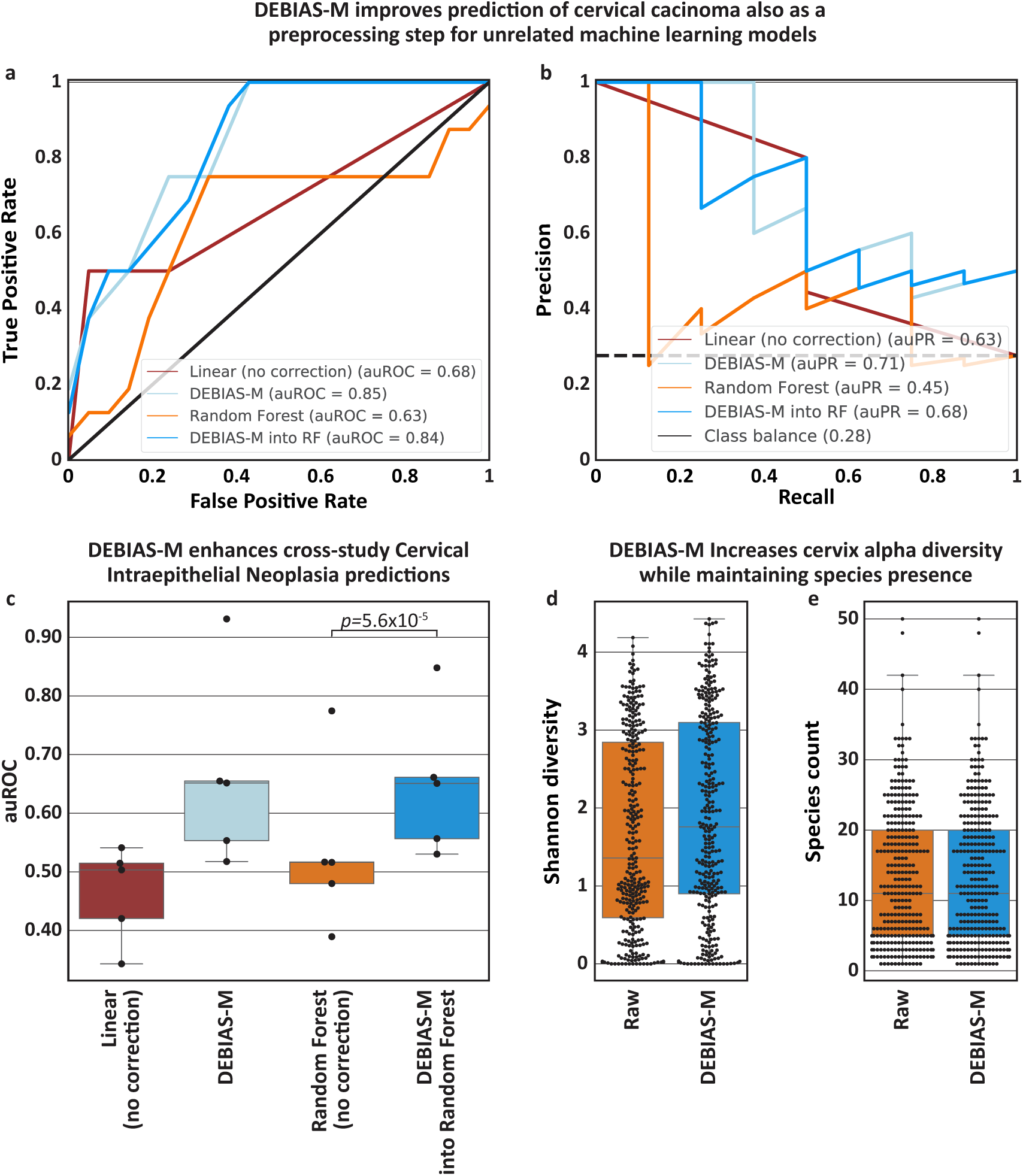
DEBIAS-M improves cross-study prediction of melanoma immunotherapy response. ROC plots (**a)** and precision-recall plots (**b**) of cross-cohort predictions of cervical carcinoma for linear and random forest models trained on cervical microbiome data, evaluating DEBIAS-M as a pre-processing step. **c,** Box and swarm plots (Box, IQR; line, median; whiskers, nearest point to 1.5*IQR) of auROCs, each evaluating the generalization performance of linear and random forest models predicting cervical intraepithelial neoplasia from cervical microbiome data to a held-out study. DEBIAS-M is used as a pre-processing step (N=29-82 samples per study). **d,** Shannon diversities of both the raw relative abundance of cervical microbiome samples and of the same samples after processing with DEBIAS-M. DEBIAS-M increases the diversity of these samples (Wilcoxon *p<*10^−20^). **e,** Count of species presence for the same samples. By design, DEBIAS-M does not increase the number of observed samples.

We next examined a challenging scenario, and replicated a cross-study analysis of predicting immunotherapy response (12-month progression-free survival) from the gut microbiome of patients with melanoma, in which the original authors concluded that there is limited generalizability of microbiome-based prediction across cohorts^58^. Analyzing data from six cohorts^58–60^ from the United Kingdom, the Netherlands, and the United States, we observed a small but consistent improvement in all cohorts analyzed when adding DEBIAS-M as a preprocessing step to the analysis pipeline implemented in the original study, and a strong improvement in the cohort of Peters et al.^60^ (**Fig. S6**). As in the cervical neoplasia analysis, use of random forest models did not improve on linear models (**Fig. S6**). Overall, these results demonstrate the utility of DEBIAS-M in realistic applications of cross-study machine learning models.

## Discussion

Sequencing-based measurements of microbial communities have demonstrated substantial variability between profiling approaches, labs, and even batches of the same study, making it challenging to replicate results across studies. To address these issues, we developed DEBIAS-M, a method for correction of microbiome processing bias. DEBIAS-M offers three notable benefits: First, it is based on a specific theoretical framework that stipulates taxon- and protocol-specific multiplicative biases^22^, and, as a result, infers bias-correction factors that are interpretable, robust, and consistent. Second, its design facilitates the development of generalizable machine-learning models, as it is able to operate without using the outcome labels of a test dataset. Third, in extensive benchmarks across more than 30 different studies, we compared DEBIAS-M to commonly used batch-correction methods and demonstrated that it consistently improved the ability of microbiome-based prediction models to predict phenotypes on held-out studies in a diverse range of clinical settings, including colorectal cancer, cervical neoplasia, and HIV, and using both metagenomics and 16S rRNA amplicon sequencing data. DEBIAS-M is available as an open-source package at https://github.com/korem-lab/DEBIAS-M.

The multiplicative processing bias framework introduced by McLaren et al.^22^ implies that even the same set of samples processed differently (e.g., using different extraction kits) might yield different microbial abundances and different associations with phenotypes (c.f. Fig. 2 in ref. ^22^ and **Fig. 1a**). This is even more likely when samples originate in different studies, with additional technical (e.g., storage time) and non-technical (diet, genetics) factors being considered, underlining the challenge of bias and batch effects in microbiome studies. In our results, these challenging differences combined with compositional confounders manifest in models that perform worse than chance when evaluated across studies, seen as auROCs<0.5 (e.g., **Figs. 2,6**).

DEBIAS-M makes two important operational assumptions. The first is that samples from different batches should generally be similar. This assumption is inherent to many batch-correction methods, and of course would not hold in some scenarios, such as joint analysis of microbiome data from different body sites. The second assumption is that there exists some association between the ground-truth (unbiased) microbiome data and the phenotype used by DEBIAS-M, which is weakened by processing bias, and therefore can be improved via bias correction. We thus expect DEBIAS-M to work better with informative phenotypes. The combination of the two assumptions ensures that DEBIAS-M does not overfit to either the cross-batch similarity or the available phenotype labels. We note, however, that the second assumption limits DEBIAS-M’s utility to certain large-scale efforts aimed at estimating measurement variability^19,61^, which typically include a small number of samples and lack an informative phenotype.

While previously developed batch-correction methods such as ComBat^29^, voom-SNM^30,31^, and ConQuR^23^ may reduce batch effects, they are not grounded in a framework that relates them to specific bias-generating processes. They might therefore produce transformations that are hard to interpret, such as introducing values to very sparse features^34,62^, and in many cases their output should likely be interpreted as a global transformation over the data rather than an attempt to quantify the abundances of specific microbes. Additionally, such methods can potentially capture and remove signals that are unrelated to processing bias, a risk that becomes more substantial when outcome labels of the test set are provided to the method. In contrast, we demonstrate that bias-correction factors inferred by DEBIAS-M are linked to biological properties of the experimental processing pipeline, and are therefore interpretable. We posit that this is due to the restrictions imposed on the correction performed by DEBIAS-M, limiting it to per-taxa multiplicative bias. While there likely are processes driving differences between batches that are not captured by multiplicative per-taxon biases, our results suggest that more flexible batch correction using contemporary methodology does not lead to improvement over the more restrictive approach of DEBIAS-M. Nevertheless, there might be specific scenarios in which a combination of approaches may be useful.

Another major difference between DEBIAS-M and currently existing methods is its suitability for developing and validating models on unseen data. Several microbiome batch-correction methods require outcome variables for all data, even for held-out test sets, leading to overfitting, “leakage” of information into the training data of the model and invalidating tests for generalization^47^. Other methods are able to perform batch correction without the use of outcome data, but with the consideration of features (microbial abundance data) from the entire dataset. While, contrary to others^47^, we do not believe this necessarily constitutes information leakage, it does limit the translation of these models. Conversely, DEBIAS-M can handle missing data, and offers substantial flexibility with respect to available microbial and outcomes data. We demonstrate that it performs similarly both when the microbial data for held-out test sets is made available and when it is kept unseen.

Importantly, our results do not attempt to identify experimental procedures that are “better” than the rest. Furthermore, observing a collection of datasets spanning different experimental protocols would help ensure that DEBIAS-M does not converge towards a microbiome representation overly biased by, for example, one particular extraction kit. However, our results showing lower variance in bias correction factors for manual processing suggest a benefit for this approach, which should be balanced with consideration of cost and practicality. We do note that our simulation results indicate that DEBIAS-M operates better with more samples per batch, with samples that represent a similar ecosystem (e.g., vaginal microbiome), and with informative phenotypes. These could serve as consideration for design of future studies, in addition to obtaining orthogonal measurements (i.e., qPCR, dilution series), which may be used to estimate bias directly^22,63^.

As consideration of multiplicative bias shows promise for microbiome data analysis, future work could investigate ways with which to assign a bias-correction factor to each processing step, as opposed to an entire study as was done here. This could facilitate the incorporation of bacterial metadata, such as Gram status, into the learning framework. In addition, positive controls and dilution series can also be incorporated by future studies as means to evaluate specific biases^63^. It is also possible that DEBIAS-M could be combined with probabilistic microbiome processing models, such as SCRuB^64^, with the intention of using as much information as possible during each of these processing components. Lastly, there are other measurements outside of the microbiome field that are susceptible to processing biases (technical or non-technical), such as transcriptomics or metabolomics, and there may be an opportunity to modify DEBIAS-M for such scenarios.

## Methods

### The DEBIAS-M generative model: processing bias

Consider a matrix *X* ∈ ℝ*^n^*^×^*^m^* representing the number of reads (or relative abundances) originating in one of *m* taxa for each of *n* samples; a vector *A* ∈ ℤ^+*n*^, where each *A_i_* ∈ {1,…, *B*} denotes the batch sample *i* originates from, with *B* total batches; and a phenotype vector Y ∈ {0,1}*^n^* that describes some information of interest that is expected to be associated with *X*.

The reads observed in each sample *X_i_* are the result of some experimental process (e.g., DNA extraction, 16S rRNA gene amplification, and sequencing). This process attempts to measure the underlying “true” composition of *m* taxa in that sample, which we denote Γ*_i_* ℝ*^m^*. Γ*_i_* represents relative abundances, such that all Γ*_ij_* ≥ 0 and Σ*_j_* Γ*_ij_* = 1. As shown by McLaren et al.^22^, for each batch *b* ∈ {1,…, *B*} and taxon *j* there exists a specific multiplicative bias term that can increase or decrease the likelihood of Γ*_ij_* to be observed in the downstream *X_i_*. These biases can be due to interbacterial differences in DNA yield, gene copies, PCR primers, extraction protocols, and so on. Since the biases inflicted by each experimental processing stage are all assumed to be multiplicative, they can be aggregated into a single bias factor per microbe within every batch^22^. To capture this phenomenon, we propose one weighting parameter *W_Aij_* >0 for each taxon-batch pair, where each *W_Aij_* represents the multiplicative bias that protocol *b* had on taxon *j*. Thus, we draw each observed sample *X_i_* from a multinomial mixture 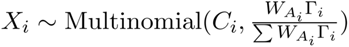, where 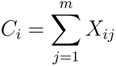 represents the total number of reads observed in sample *X_i_*.

Thus, given parameters *W* ∈ ℝ*^B^*^×*m*^ and Γ*_ij_*, the probability of an observed dataset *X* is:

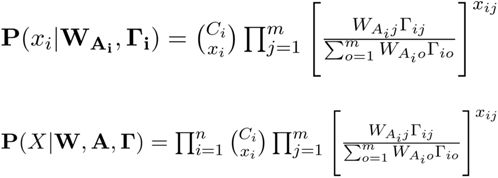

### The DEBIAS-M generative model: phenotypic associations

Next, we assume that there exists an association between the underlying Γ and the phenotype *Y*. While DEBIAS-M could model this association in various way, here we chose to use logistic regression, with a single set of linear weights *L* ∈ ℝ*^m^*, such that *Y*|Γ,*L*∼*Bernoulli*(*Y*,*θ*(*L*Γ)), where *θ* is the sigmoid function. We assume that in the absence of processing bias, the association between Γ and *Y* is protocol- and batch-independent, and therefore the weights *L* should be identical across all batches. Given this, the probability of observing a set of labels *Y* given Γ and *L* is:

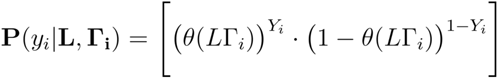

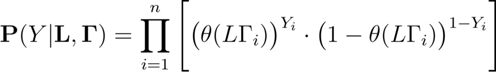

By combining the two components together, we can link the observed read counts *X* to the phenotype *Y* through the processing biases *W* and linear weights *L*:

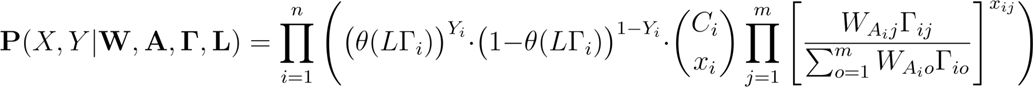

### Cross-batch similarity

We next make an additional assumption, that batches of the same sample types in similar contexts should be similar, as is standard for established batch effect correction methods^23,24,26,33^. This would generally apply to microbiome data collected from the same type of environment (e.g. human gut from studies sharing the same patient exclusion criteria). We express this assumption by comparing the L2 distance between the pairwise average of the Γ inferred for each batch. Thus, we introduce *μ* ∈ ℝ*^B^*^×^*^m^* = *μ*(Γ,*A*), the average of each batch:

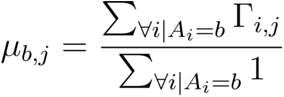

We assume there exists an underlying probability in which batch means 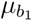 and 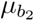 are equivalent. Then, assuming that each *μ* follows a multivariate Gaussian distribution which we set with a covariance matrix of Σ=*αI*, where *I* is the identity matrix and *α* is a scaling hyperparameter, such that all *μ_b_* ∼ 𝒩(*μ*’, *αI*). We offer a simplification to directly optimize the pairwise L2 penalty in logspace, which would amount to the following expression if we consider each pairwise batch averages as a variable/mean-parameter combination:

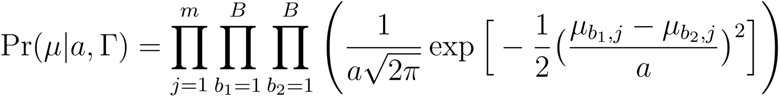

We use this approach to produce an L2 penalty, but note that a similar approach could be used to produce an L1 penalty.

Adding this cross-batch similarity term into the generative model yields:

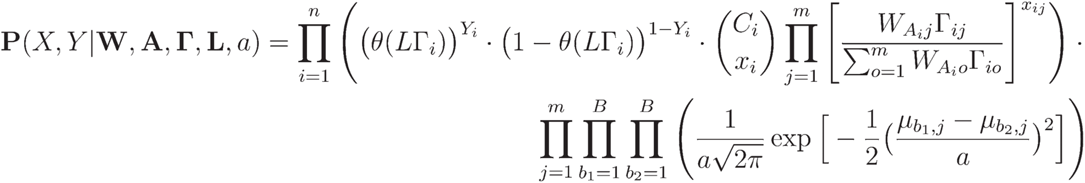

### Modeling

Given observations for *X*, *Y*, and their corresponding batches *A*, we aim to infer the parameters *W* and *L* of the generative framework. We learn these parameters through stochastic gradient descent, after initializing *W* to **1**. For our model, we use an inverse of *W*, *W*’, such that, for any *X_i_*:

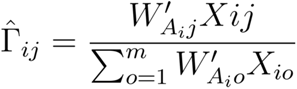

Whereas *W* is the processing bias, *W*’ could be interpreted as a “bias-correction factor”. For simplicity, we maintain the notation of *W* below. We then use this inferred Γ in **P**(*X*,*Y*|**W**, **A**, **Γ**, **L**), which we assess in log-space:

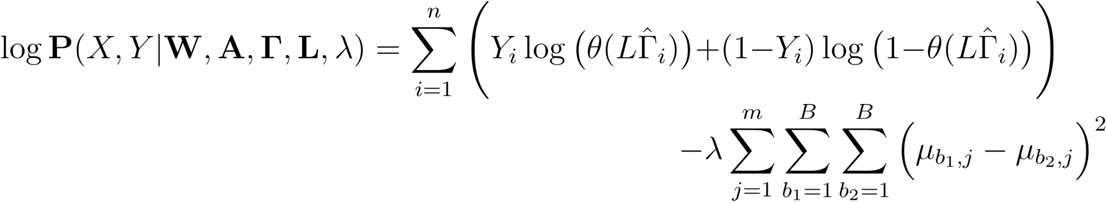

Where we absorb the *α* hyperparameter into the hyperparameter *λ*.

Using this expression for log **P**(*X*,*Y*|**W**, **A**, **Γ**, **L**, λ), we iteratively backpropagate all the way through the Γ*_ij_* estimate to identify which *W*,*L* parameters maximize the log-likelihood. Importantly, we note that while we include a modeling term that accounts for a cross-entropy loss associated with a phenotype of interest *Y*, this framework can also account for observed samples in the Γ and *μ* terms for which the phenotype of interest is not yet known. Therefore, this approach allows us to correct for batch effects while maintaining proper train/validation/test splits, making use of batches in which either partial labels or no labels are available. While we omitted traditional regularization methods like lasso and ridge penalties from the above equations, we note that they could be applied to both the *L* and the log of the *W* weights.

### Implementation of DEBIAS-M

DEBIAS-M is implemented in pytorch-lightning^65^, using the adam^66^ optimizer with a learning rate of 0.005 and otherwise default parameters, run with at least 25 epochs. Before the DEBIAS-M optimization, the linear weights *L* are initialized using the unregularized scikit-learn^67^ LogisticRegression model trained using the uncorrected data. The log2(W) are stored as free parameters, thus ensuring that all the *W* are non-negative. The Γ are renormalized to relative abundance during the DEBIAS-M forward step. During training, sample batches are selected during the predictive modeling component, while all samples are incorporated into the cross-batch difference measurements. While the hyperparameter *λ* can be tuned via cross-validation, we found that DEBIAS-M performs well when it is weighted as a function of the number of features and possible pairwise batch comparisons. We therefore define 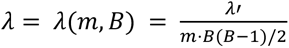 and set *λ*′ = 10^4^ by default. We empirically found that 10^4^ is an effective scaling value; this expression was used as our *λ*′ weight for all analyses.

Unless noted otherwise, we allow samples from both the training and test set to be used in the cross-batch similarity terms, while only samples from the training set are used in the cross-entropy loss and predictive weights. This implementation most closely mirrors that of standard batch-correction models, in which the input data (i.e., microbiome data) from all batches are observed. In our ‘adaptation’ benchmark (**Fig. S1d, S3**), however, the *L* and *W* weights are optimized only for the training set, without observing the test set. After those weights are frozen, the *W* weights for the test set are adapted to optimize its similarity to the training set, while the predictive model layer (i.e., the *L* weights) do not change during this process. Those predictive parameters are then applied to the test set exactly as they were learned when only observing the training set. This setting is more conservative, and keeps a full separation between the input data of the train and test sets.

All implementations and per-model training of DEBIAS-M used in this work required less than one minute of runtime on a standard laptop.

### Microbiome data acquisition and processing

All datasets used in this work were publicly available at the time of analysis. We obtained the HIV data^35^ from Synapse (https://www.synapse.org/#!Synapse:syn18406854), using the ‘taxonomic_assignments/insight.merged_otus.txt’ file with data processed using Resphera Insight^35^. We obtained the cervical neoplasia data from the repository available for each study (**Table S2**), and processed it with DADA2^68^. We used any indication of cervical intraepithelial neoplasia as phenotype in analysis. All studies in this dataset were used for training subsets, but only studies that had subjects with both phenotype labels were included in evaluation. The colorectal cancer data was obtained from the R curatedMetagenomicData package^69^, which provides species-level relative abundance data processed by MetaPhlAn3. Bacterial metadata was obtained from https://gold-ws.jgi.doe.gov/.

### Benchmarking of batch-correction methods

For all benchmarks, we ran batch-correction methods on the raw read counts after adding 1 to all values. We ran: (1) ComBat using the ‘ComBat_seq’ function; (2) ConQuR using the ‘ConQuR’ function; and (3) voom-SNM using code made available by Poore et al. (file ‘Plasma-Voom-SNM-Normalize-Age-and-Sex.R’)^33^. As ConQuR and Voom-SNM require a covariate variable and do not withhold that variable from the test set, we used gender for the HIV and colorectal cancer studies and age > median for the cervical neoplasia datasets. The outputs of Voom-SNM, ConQuR, and Combat, along with unmodified (raw) relative abundances, were assessed using logistic regression (scikit-learn^70^) with no penalty. For DEBIAS-M, the correction and prediction are implemented simultaneously through a similarly unregularized linear layer, without considering any metadata except for the outcome label of the training data. Of note, such separation between training and testing data is not available for other batch-correction methods.

We implemented a cross-study validation pipeline, in which we trained a model on data from all but one study, and evaluated the predictive performance of the model on the held-out study, such that in the boxplots in **Figs. 2a-c, 6c, S2, S3c, S4a-c, S6** each “dot” represents that held-out study. To compare the performance of multiple classifiers on the same prediction benchmark (e.g. HIV classification following DEBIAS-M and an alternative method) we used Fisher’s multiple comparison to combine DeLong tests performed on each pairwise comparison of performance on the same held-out study.

For the prediction benchmarks in **Fig. 6**, we ran the same general predictive pipeline for cervical carcinoma and cervical intraepithelial neoplasia, but rather than using a logistic regression model, we used a random forest model. For this model, we tuned the max depth and max features hyperparameters using cross-validation on 3 folds (nested within training data). For the prediction benchmark in **Fig. S6**, we considered both the same random forest tuning pipeline, and an L1 logistic regression model of the relative abundance data in logspace with a pseudocount of 10^−4^, as in the original analysis^58^. In both cases, we also tuned the hyperparameters of DEBIAS-M (with nested cross-validation): the learning rate, L2 regularization strength, and *λ*′ (using 10^3^, 10^4^, and 10^5^).

### DEBIAS-M in logspace

As it is common and effective to analyze microbiome datasets in log-space, we also provide a version of DEBIAS-M tailored to this feature space. As bias terms are multiplicative in count and relative abundance space, they are additive in logspace. We therefore refer to this version of DEBIAS-M as ’log-additive DEBIAS-M’ (used in **Figs. S2, S4**).

In clr-transformed space, the unbiased Γ̂*_ij_* term can be represented with the following expression, which includes normalization that mimics the same renormalization process in relative abundance space:

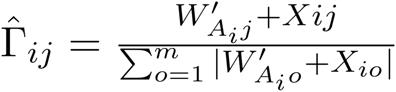

We note that different logarithmic transformations would likely require different normalization terms. Following this log-additive adjustment, the rest of the DEBIAS-M optimization applies to Γ*_ij_*.

However, we modify the *λ*’ hyperparameter to 10^-^, to account for the larger range of values in the clr-transformed data. In benchmarks involving log-additive DEBIAS-M, we used similar LogisticRegression models, but transformed the count matrices produced by ConQuR and ComBat using the centered-log-ratio (clr) transform. We omitted Voom-SNM from this analysis because its output is not in count space.

### Simulation of synthetic processing biases

To evaluate the robustness of the DEBIAS-M model across different scenarios, we used data-driven simulations to measure the bias-correction against a known ground truth. To this end, we used SparseDOSSA2^49^ to generate simulated data, which we trained on the vaginal microbiome data provided in Dataset S2 of DiGiulio et al.^50^ using default parameters. With this trained model, we generated 25 synthetic datasets of 1,000 features and 384 samples, from which we created simulations of batch effects. We varied: 1) the number of batches, using 2, 3, or 4 batches; 2) the numbers of samples per batch, using 24, 48, 72, or 96 samples; and 3) the number of features, using 100 or 1000 features.

To simulate phenotype labels, we randomly generated linear weights for each feature, drawn i.i.d from a Gaussian distribution of mean 0 and standard deviation of 2. Multiplying these simulated linear weights by each sample yielded a score for each sample, which is by construction perfectly associated with the simulated microbiome. We then modulated the strength of this association by adding noise drawn i.i.d from a Gaussian distribution with a mean of zero and standard deviation of 0.1, 1, or 10.

These were used to generate the final phenotype label – with “strong”, “moderate”, and “weak” associations, respectively – which were binarized using the median as a threshold. The final labels corresponded to average permanova R^2^ with the simulated microbiome data data of 0.086 (median *p*≤10^−5^), 0.041 (median *p*≤10^−5^), and 0.007 (median *p=*0.23) for “strong”, “moderate”, and “weak”, respectively.

To simulate bias, we drew log2 bias factors for each study-feature combination from i.i.d Gaussian distributions of mean 0 and standard deviation of 2. The exponents of these factors were then multiplied by each sample from the corresponding study, before being proportionally renormalized to simulate sequencing depths of 10^3^, 10^4^, or 10^5^. For every experiment, all biased samples and labels were provided to DEBIAS-M, and the output samples were then compared against the known simulated ground-truth via jensen-shannon divergence (JSD). The same comparison was also made for the uncorrected samples. The median JSD for each experiment was then recorded, for a total of 25 points per box in **Fig. 3**. For all plots, the default settings used for parameters other than the one being investigated in that particular panel were 4 studies, 96 samples per study, 1000 features, low phenotype noise, and read depth of 10^5^.

### Inference of DEBIAS-M bias-correction factors

To investigate the bias-correction factors inferred by DEBIAS-M, we utilized the collection of HIV datasets, which included studies with a wide range of experimental designs. We began by implementing DEBIAS-M using our standard train-test split, but running it twice per validation batch, with each iteration observing a randomly selected half of the batch. The resulting inferred bias correction factors for OTUs that were present in both semi-batches were compared in aggregate across all studies (**Fig. 4a**). Next, we evaluated a DEBIAS-M model that included the samples and labels of all batches. The resulting bias-correction factors were analyzed via adonis^51^ and principal component analysis. While in the analysis in **Fig. 4** bias-correction factors for non-detected taxa were kept at 1, in **Fig. S5** we instead imputed them to the largest observed bias-correction factor, implying an assumption that maximal bias against the taxa caused it to go undetected. We also evaluated the detection (presence/absence) of each taxa in the HIV studies directly, and evaluated these differences via adonis and agglomerative clustering algorithm, using the Manhattan distance metric and the complete linkage method (using the R stats^71^ package). Gram status and 16S copy number were obtained from the GOLD database^72^.

### Analysis of multi-task DEBIAS-M

The benchmark for prediction of metabolite abundance used the metabolite data and metadata from Table S1 in Kindschuh et al.^3^, and the microbiome data from Supplementary Data 2 in Elovitz et al.^55^, which was processed in two batches. For each of the 509 metabolites with multiple unique observations in both microbiome batches, we evaluated a prediction task in which we aimed to predict if the level of a metabolite in a particular sample was greater than the median of that metabolite across the entire dataset. We used the larger batch as training data and the smaller one as a test set.

We ran the linear baselines for the raw, Combat, and ConQuR datasets as before. Additionally, we ran a multitask version of the DEBIAS-M model, in which a collection of multiple *L* parameters, one for each metabolite, were simultaneously learned alongside a single set of *W* weights (Fig. 5a). As an additional benchmark, we applied MelonnPan^57^ to the same prediction task, in which we trained it on the larger batch and made predictions on the smaller batch. We then assessed MelonnPan’s predictions against the same classification framework to obtain auROCs for all of the 120 metabolites for which MelonnPan provided predictions, and compared the output of multitask for the same set of metabolites (Fig. 5c).

## Code availability

DEBIAS-M is available from https://github.com/korem-lab/DEBIAS-M. Code used to generate all analyses and plots can be found at https://github.com/korem-lab/v1-DEBIAS-M-Analysis/.

## Data availability

All datasets analyzed in this study are publicly available. The HIV dataset is available from Synapse (https://www.synapse.org/#!Synapse:syn18406854). The colorectal cancer and melanoma immunotherapy datasets are available through the R curatedMetagenomicData package^69^. The cervical neoplasia dataset was compiled from data provided with each publication, with information detailed in **Table S2**.

## Acknowledgements

We thank members of the Korem group and Gregory D. Sepich-Poore for useful discussions, and reviewers from RECOMB2024 for insightful comments. We thank all authors and participants involved in the generation of data used in this study. This work was supported by the Program for Mathematical Genomics at Columbia University (T.K.), R01HD106017 (T.K.), and T15LM007079 (G.I.A.).

## Author contributions

G.I.A. and T.K. conceived and designed the study, designed analyses, interpreted the results, and wrote the manuscript with input from A.B.K., H.P., J.B. and A.-C.U. G.I.A. conceived and wrote DEBIAS-M and conducted all analyses. J.B., A.B.K and H.P. acquired, harmonized, and processed data.

J.B. and A.B.K. assisted with analyses. T.K. supervised the study.

## Supplementary figures

**Figure S1.**
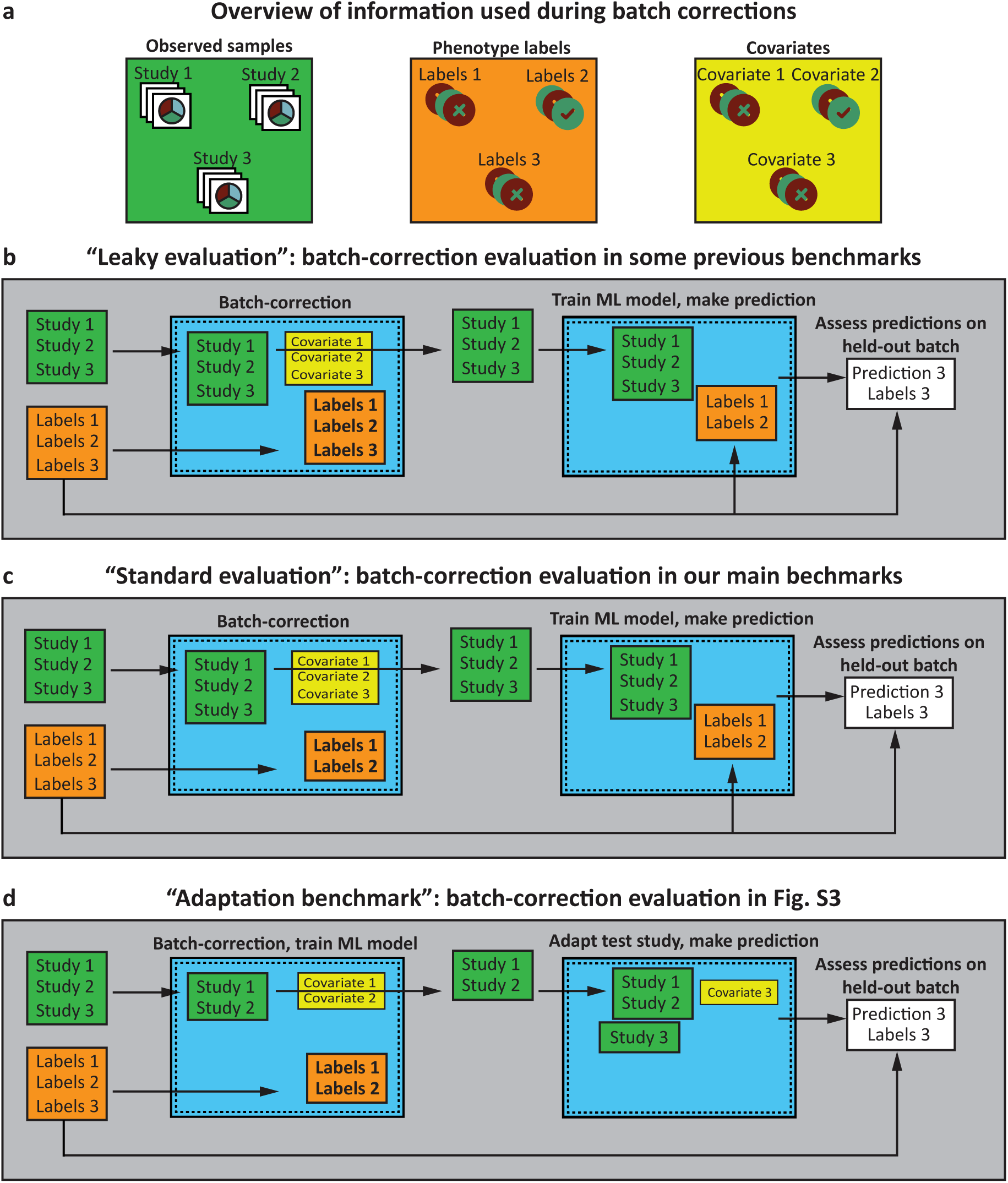
Overview of information used in various microbiome batch-correction prediction benchmarks. **a,** Description of information that is typically incorporated in microbiome batch correction, which is 1) the samples themselves; 2) the labels to be predicted in downstream modeling; and 3) other covariates. **b,** An approach that has been used in some previous benchmarks, in which the labels of the test set are used during batch correction itself. This risks “information leakage”, and is used in this work only in **Fig. S2. c**, The primary batch-correction evaluation strategy used in this work for DEBIAS-M, in which the samples and covariates from all studies are used during batch correction, but only the labels from the training set are used during batch correction or model training. **d**, The batch-correction strategy used in our ‘adaptation’ benchmark in **Fig. S3**, in which no information from the test set is used during batch correction or model training. Once all bias-correction factors and predictive model weights are learned and fixed for the training set, bias correction is performed separately for the test set by adjusting its bias-correction factors to optimize cross-batch similarity.

**Figure S2.**
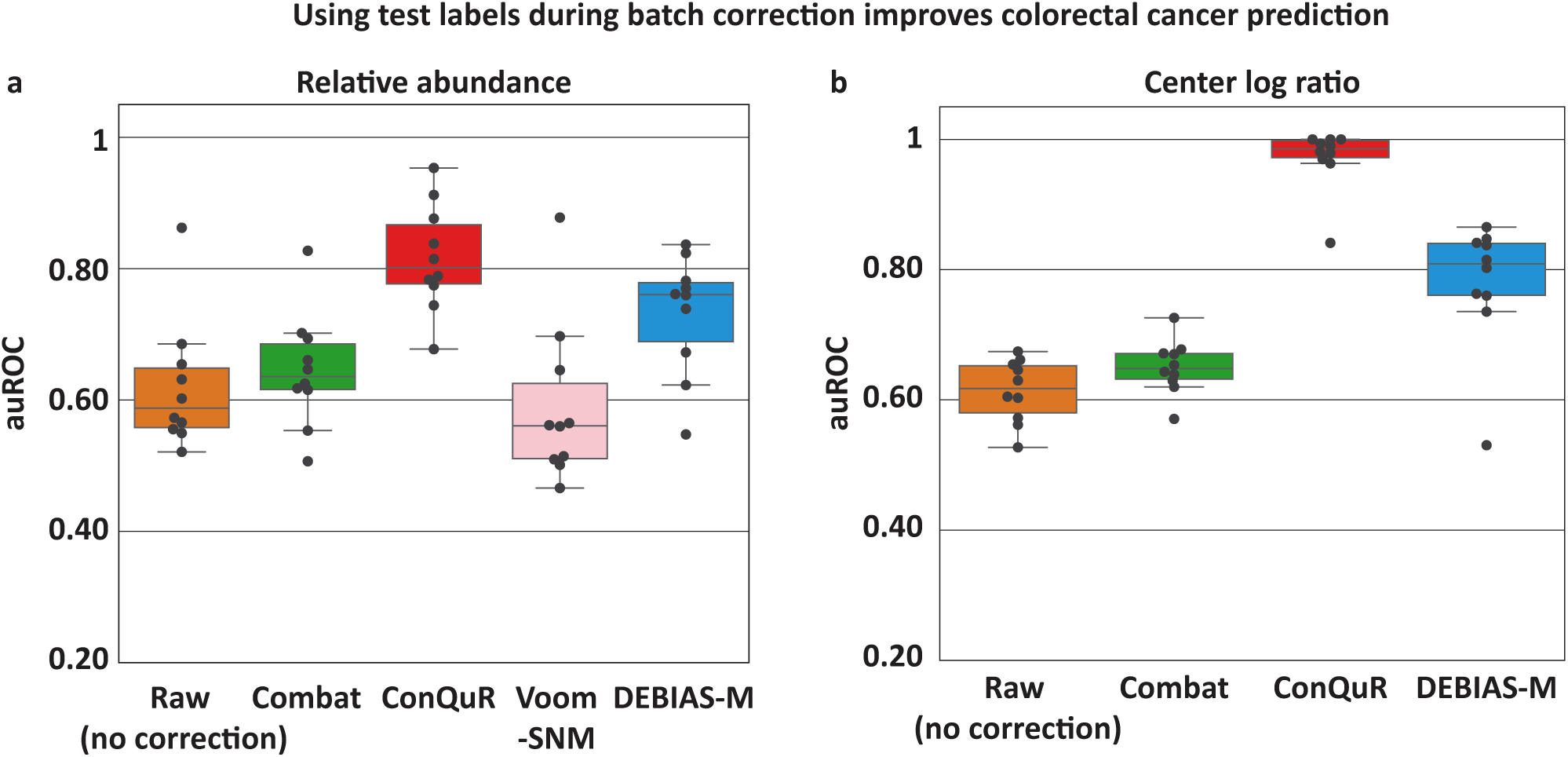
Using test set labels during batch correction can drastically increase measured predictive performance in downstream benchmarks. a-b, The same cross-study colorectal cancer prediction benchmark as in Fig. 2d, but Combat, ConQuR, and Voom-SNM were provided all colorectal cancer labels, including for the test set, during batch correction (**Fig. S1a**). The prediction accuracy (auROC) of certain methods inflated drastically beyond the results observed in the primary benchmark (Fig. 2d), highlighting potential issues with assessing a batch-correction method by measuring the ability of a downstream machine learning model to predict information used during batch correction. This trend is consistent in both relative abundance space (**a**) and center log ratio (**b**). Voom-SNM is not run for (b) as its output is neither in non-negative relative abundance nor in count space.

**Figure S3.**
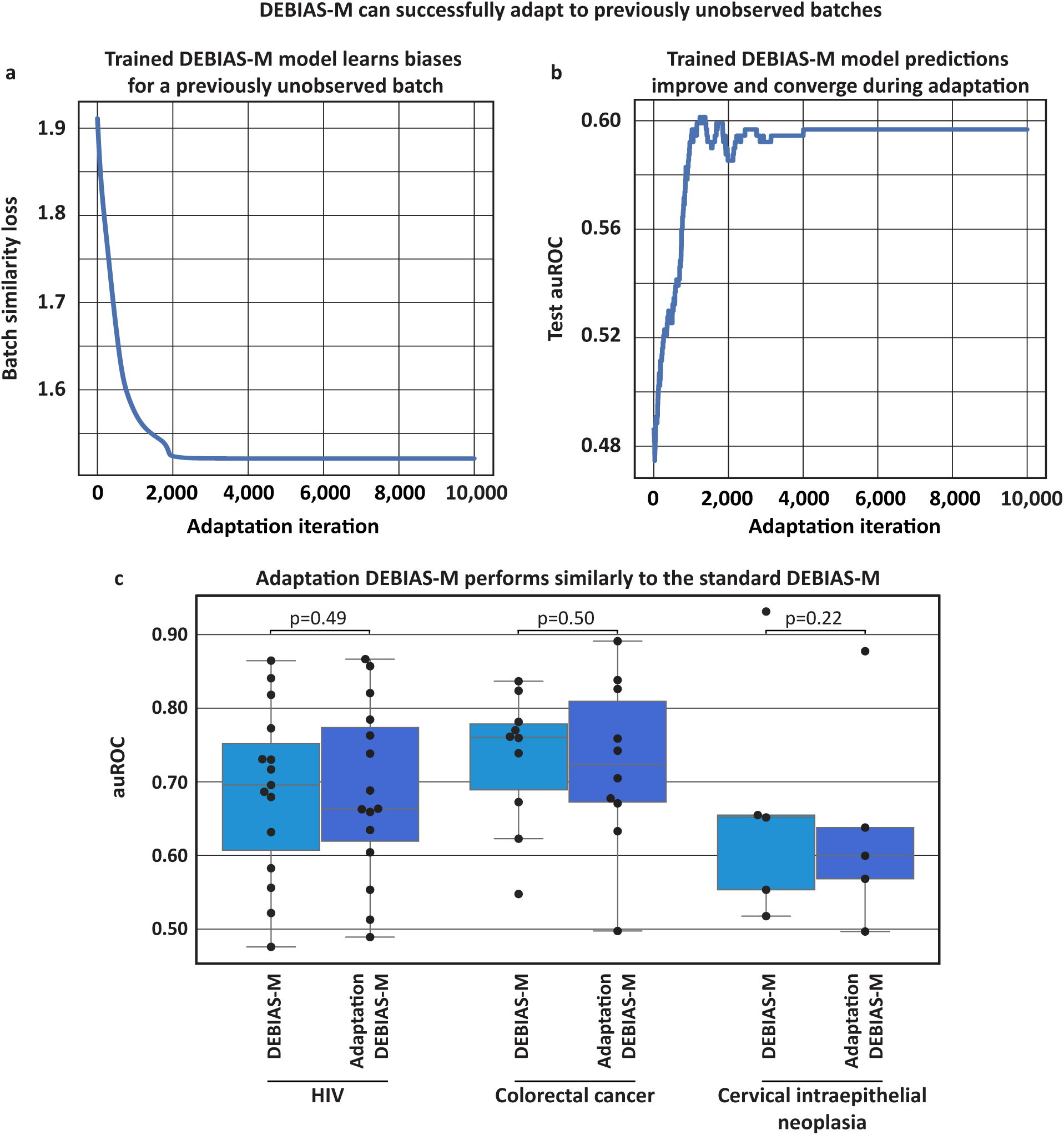
A fitted DEBIAS-M model effectively adapts to previously unobserved samples. **a,** The progression of cross-batch similarity loss as a fitted DEBIAS-M model adapts to samples from a previously unobserved study, by solely minimizing the cross-batch similarity loss. **b,** the predictive performance of the fitted DEBIAS-M model throughout the adaptation iterations. Although not directly used during the adaptation itself, the auROC of thee model’s prediction on the held out test increases as the cross-batch similarity increases.**c,** Box and swarm plots (Box, IQR; line, median; whiskers, nearest point to 1.5*IQR) comparing the performance of DEBIAS-M (fitted and evaluated using the strategy in **Fig. S1b**) with “Adaptation DEBIAS-M” (fitted and evaluated using the strategy in **Fig. S1c**) on the same benchmarks used in Fig. 2. Adaptation DEBIAS-M demonstrated equivalent predictive performance on held-out studies. *p* - one-sided Wilcoxon signed-rank test.

**Figure S4.**
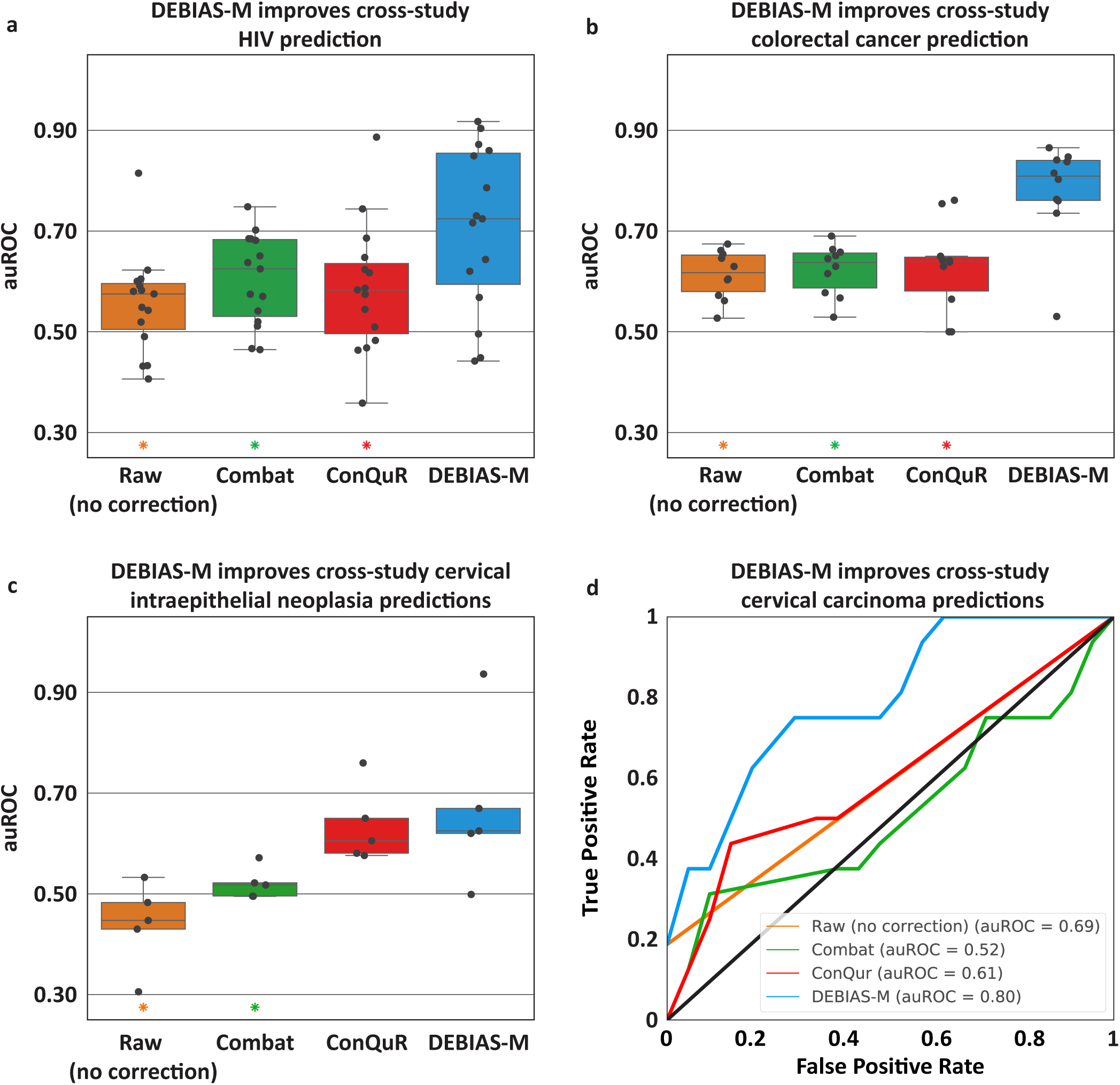
Log-additive DEBIAS-M outperforms batch-correction methods in cross-study prediction benchmarks in centered-log-ratio space. Same as Fig. 2, but comparing log-additive DEBIAS-M to batch-correction methods on clr-transformed data. Voom-SNM is not included in this benchmark as its output is not in non-negative relative abundance or count space.

**Figure S5.**
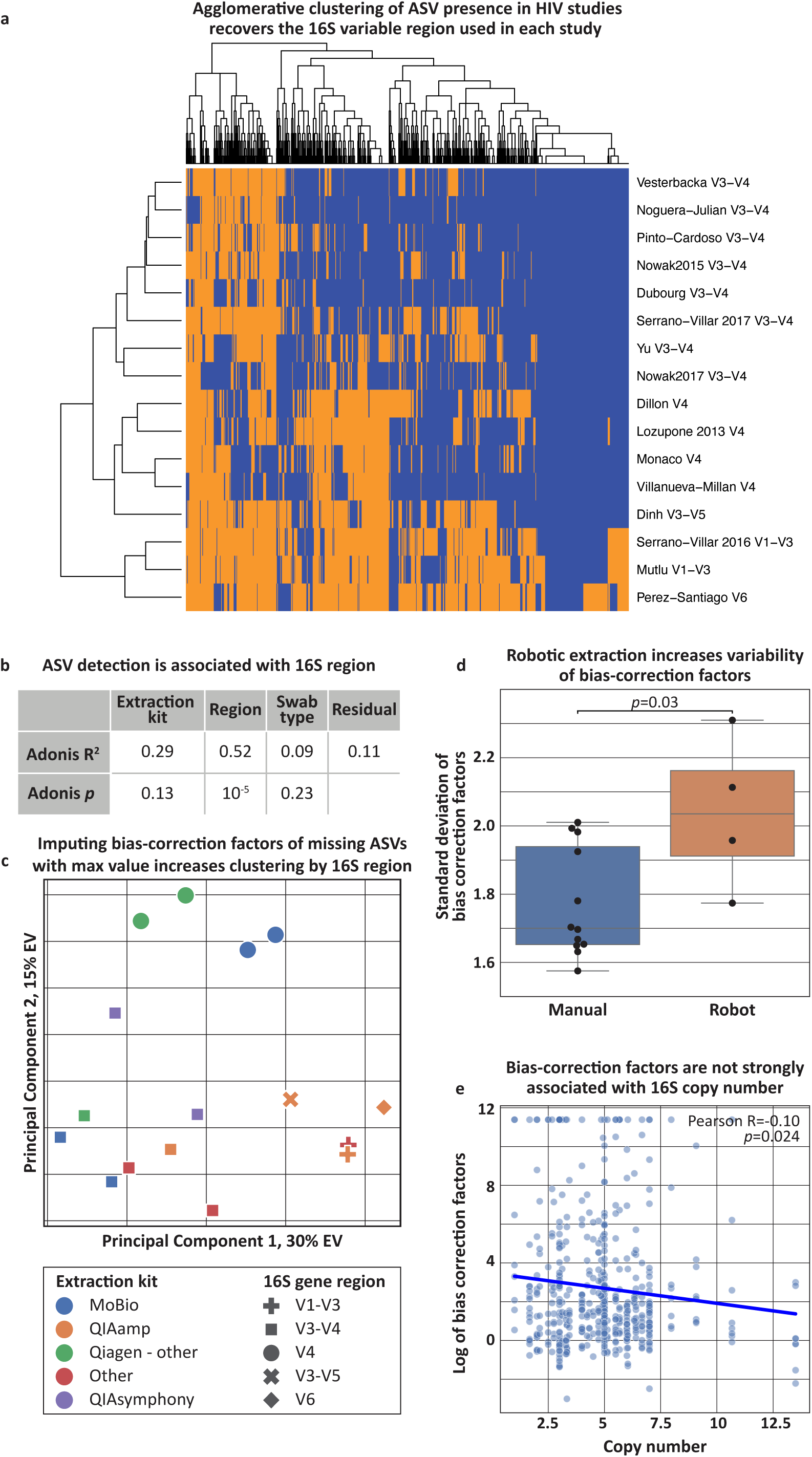
DEBIAS-M inference yields biological insights into sequencing bias. Analyses of a fitted DEBIAS-M model on the collection of HIV studies used in Fig. 2a**, 4,** with bias-correction factors for species not found in a certain study imputed to the largest observed factor across all datasets. **a,** Heatmap illustrating the presence (blue) and absence (orange) of each OTU across each of the HIV studies analyzed, displayed using agglomerative clustering (**Methods**). The OTU detection patterns of the different studies cluster according to the 16S region amplified. **b,** Adonis PERMANOVA explained variance and p values for the effect of different experimental factors (**Table S1**) on the detection (presence/absence) of each OTU across each HIV study. **c,** PCA plot of the bias-correction factors inferred by DEBIAS-M, same as Fig. 4c, but with bias-correction factors for OTUs not found in a certain study imputed to the largest observed factor across all datasets. Color represents extraction kit type and shape the 16S rRNA region used. **d,** Box and swarm plots (Box, IQR; line, median; whiskers, nearest point to 1.5*IQR) showing the standard deviation of bias-correction factors, comparing studies with manual and robotic processing. *p*, Mann-Whitney U test. **e,** Scatterplot showing the bias-correction factors inferred by DEBIAS-M plotted versus the 16S copy number of the same species.

**Figure S6.**
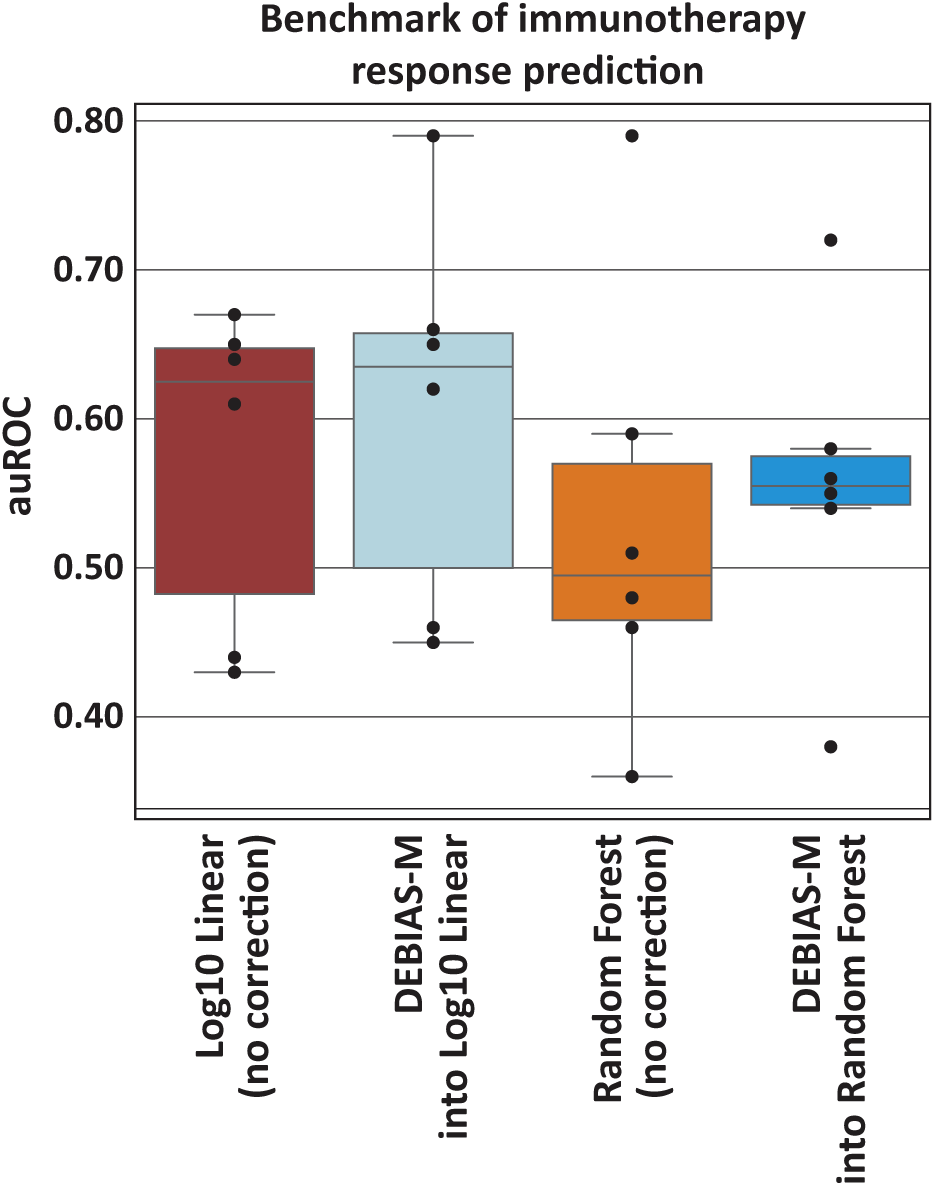
DEBIAS-M improves cross-study prediction of melanoma immunotherapy response. Box and swarm plots (Box, IQR; line, median; whiskers, nearest point to 1.5*IQR) of auROCs, each evaluating the generalization performance models using gut microbiome data to predict immunotherapy response in melanoma patients (defined as 12-month progression-free survival^58,73^). Each auROC is calculated on a held-out study. ‘Log10 Linear” denotes the pipeline used by Lee et al.^58^, with DEBIAS-M used as a pre-processing step, Preprocessing with DEBIAS-M shows a consistent albeit small improvement across all studies, with a particularly strong effect for one study.

